# Principles for simultaneous measurement of excitatory and inhibitory conductances of single cells in a single trial

**DOI:** 10.1101/690719

**Authors:** Ilan Lampl

**Affiliations:** Department of Neurobiology, Weizmann Institute of Science, Rehovot, Israel

## Abstract

Neuronal activity is determined by the interplay between excitatory and inhibitory inputs of individual cells. Whether or not these inputs covary over time or between repeated stimuli remains unclear due to the lack of experimental methods for measuring both inputs at the same time. Current methods for conductance measurement are based on repeatedly stimulating neurons while holding their membrane potential at different levels so to reveal either excitation or inhibition, which can only provide the averaged relationship between the two. Here we develop a new framework for simultaneously measuring both the excitatory and inhibitory inputs of single cells in a single trial under current clamp. This method is based on theoretical analysis of passive circuits and can be practically achieved by injecting a high frequency sinusoidal current and then analysing the data using an optimization algorithm. We use simulations to demonstrate the ability of this approach to reveal the excitatory and inhibitory inputs of point neurons, in which we mimic adapting sensory inputs as well as an asynchronous balanced state.

## Introduction

Neurons throughout the nervous system typically receive both excitatory and inhibitory synaptic inputs, and the relationship between the two determines the neurons’ subthreshold potential and firing times. As such, studying this relationship has been crucial to solving fundamental questions in neuroscience, both in health and disease. For example, its role in determining the selectivity of neurons to sensory stimulation has been studied extensively in different modalities in-vivo using either voltage clamp or current clamp approaches. Disrupted relationships between these inputs was suggested to accompany many neurological diseases and in particular elliptic seizures. It is commonly believed that such seizures are accompanied and even caused by increased excitation-inhibition ratio (Schevon et al., 2012). (McCormick and Contreras, 2001; Schevon et al., 2012).

In the voltage clamp approach, the membrane potential is clamped near the reversal potential of inhibition (∼ −80 mV), to reveal excitatory synaptic currents and near 0 mV to record inhibitory currents. Voltage clamp recordings have been used in this matter to study mechanisms of feature selectivity of cortical cells belonging to various modalities ((Borg-Graham et al., 1998; Haider et al., 2010; Atallah et al., 2012; Adesnik, 2017); (Wu et al., 2008; Zhou et al., 2014; Moore et al., 2018). Some studies used such recorded currents to compute changes in excitatory and inhibitory conductances due to sensory stimulation ((Wehr and Zador, 2003; Tan et al., 2004; Li et al., 2012). Current clamp recordings have been used in a similar manner to reveal excitatory and inhibitory conductances, by injecting constant positive or negative currents which bring the membrane potential near the reversal potential of these two inputs ((Anderson et al., 2000; Priebe and Ferster, 2005; Wilent and Contreras, 2005; Heiss et al., 2008; Scholl et al., 2013). The two approaches share several similarities. In both cases excitation and inhibition are recorded in different trials and conductances are estimated with the membrane potential equation (Eq. 1 below) while fitting the averaged data. Usually, many trials are collected and averaged together for each holding potential. Hence, these methods provide only an average picture regarding the relationships between excitation and inhibition at each stimulation condition.

To reveal the instantaneous relation between excitation and inhibition in-vivo, a different approach was developed, using the finding that the membrane potential of nearby cortical cells in intact animals is highly synchronized (Lampl et al., 1999; Poulet and Petersen, 2008). This approach consists of depolarizing one cell in order to reveal its inhibitory inputs while simultaneously hyperpolarizing a neighbouring cell to reveal its excitatory inputs. Doing this showed that excitatory and inhibitory synaptic inputs are highly correlated in anesthetized ((Okun and Lampl, 2008; Arroyo et al., 2018)) as well as in awake rodents ((Okun and Lampl, 2008; Arroyo et al., 2018). A similar approach was also used to study the degree of correlation during oscillatory neuronal activities (Adesnik, 2018). Although the instantaneous relationship between excitatory and inhibitory inputs can be revealed by this paired recording approach, the maximum inferred degree of estimated correlation between excitation and inhibition is bounded by the amount of correlation between the cells for each input, which may change across stimulation conditions or brain-state (Reimer et al., 2014; Vinck et al., 2015; Stringer et al., 2019).

Here we describe a new framework for simultaneously measuring both excitatory and inhibitory conductances under current clamp in single trials using whole cell recordings, without making any statistical assumptions about the inputs and at high temporal resolution. It is based on analysis of passive circuits and can be applied by injecting a high-frequency sinusoidal current that creates a small sinusoidal response of no more than a couple of millivolts across the membrane, therefore also allowing one to find how these inputs shape the natural membrane potential dynamics. We explain the physics behind this approach, the computations needed for the measurement and demonstrate it using numerical simulations of single cells.

## Results

We sought to develop a method that provides a way to simultaneously measure the excitatory and inhibitory conductances with high temporal resolution under current clamp. We show that for a passive point neuron (Figure 1A, see details in legend) that E&I input conductances can be revealed if we know the recorded membrane potential (Vm) and the total conductance (G) at each time point. In this simple transformation we assume that available for us are also the total capacitance and the reversal potentials of excitation and inhibition (Figure 1). This transformation is given by the membrane potential equation (Eq. 1) and its modified version (Eq. 2).

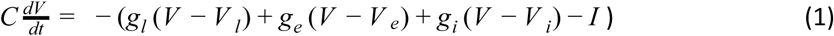

**Figure 1.**
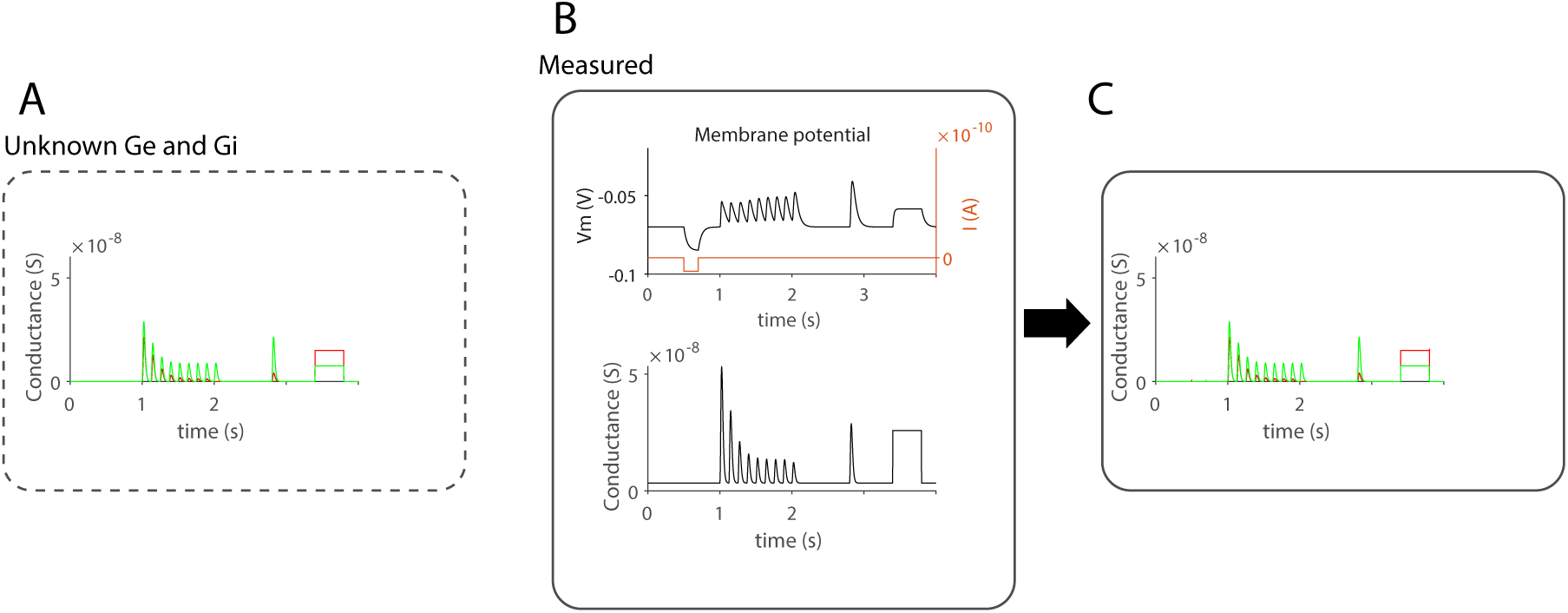
Ge and Gi can be obtained from Vm and total G. (A) Simulation of synchronized excitatory and inhibitory synaptic inputs (inhibition delayed by 4 ms after excitation), which depressed according to a mathematical description of short-term synaptic depression (STD, (Tsodyks and Markram, 1997)). These are the inputs the method aims to reveal. (B) Membrane potential simulation of a passive point neuron (*R = 300MΩ, C = 1.5*10*^*-10*^ F, Euler method, dt = 0.0005s) receiving the inputs in A, with the total conductance shown below. We assume that these two vectors are measurable. A short pulse current was injected at the early part of the trace. (C) The result of transforming of Vm, its derivative (not shown) and the total conductance into Ge and Gi using equation 1.

Replacing *V-V*_*I*_, *V-V*_*e*_, *V-V*_*i*_ with *V*^*l*^, *V*^*e*^ and V^i^, *g*_*s*_ = *g*_*i*_*+g*_*e*_, and reorganizing (1) yields:

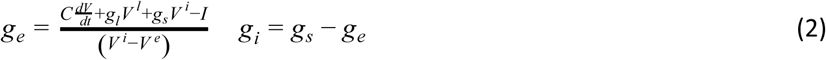

In this transformation the leak conductance is given from the baseline of the total conductance, when the membrane potential is at its resting level. Whereas the capacitance is easy to measure (from the input resistance of the cell and its time constant (*C = τ*_*m*_*/R*_*in*_)), a method for measuring the total conductance of a cell during current clamp is currently missing.

We found that under some simple conditions the total conductance of a neuron can be estimated following an injection of high-frequency sinusoidal current. This process involves several steps and depends on the recording configuration and in particular on the existence of a stable and sufficiently high series resistance (i.e. the combination of pipette and access resistance during the whole cell patch recording). Below we describe the theory behind this measurement followed by some practical steps, which we developed under simulations of point neurons.

### Impedance analysis of a simulated passive point neuron

We started by analyzing the impedance of an RC circuit in the frequency domain, mimicking the passive property of a point neuron (composed of a resistor, R (1/G, where G is the conductance of the cell) and a capacitor, C). We find that by adding a series resistor (Rs), representing the recording electrode, the absolute impedance (|*Z(f)*|) of the circuit behaves in a way that is not predicted from the cell alone (Figure 2A, B). In the absence of Rs, the impedance-frequency curves obtained for different values of cell’s conductance never intersect. However, when Rs > 0, the curves intersect so that at high frequencies, |*Z(f)*| increases rather than decreases with the conductance of the cell (Figure 2B). The total conductance of this circuit is given by Equation 3 and its absolute value by Equation 4 (w = 2*π*f).

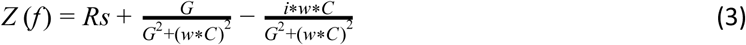

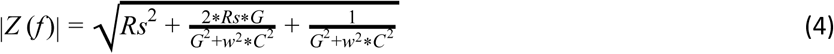

**Figure 2:**
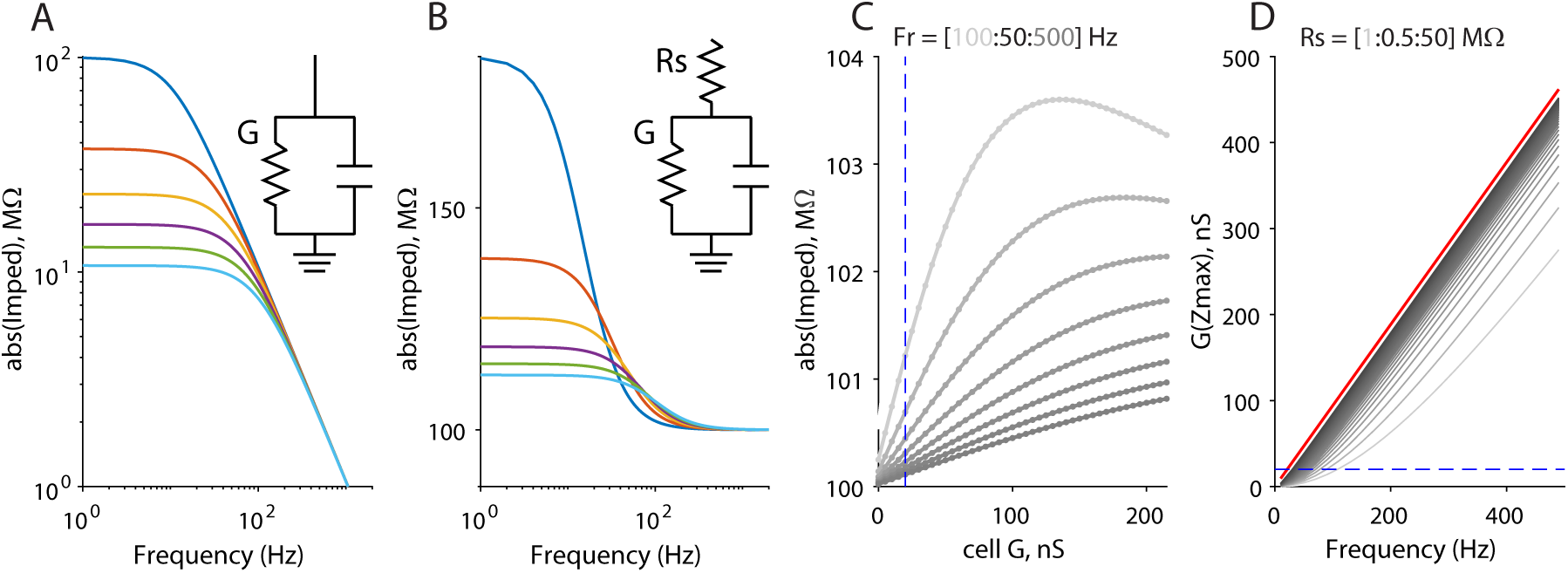
Theoretical impedance-conductance relationships for two circuits. (A) For a simple RC circuit the absolute impedance as expected is lower when cell’s conductance increases. For this circuit, impedance-frequency curves never intersect each other. (B) The impedance-frequency relationships are similar at low frequencies when a series resistor is added to the RC circuit (Rs), but for this circuit impedance-frequency curves intersect each other at higher frequencies, implying that at high frequencies, impedance increases with increasing cells’ conductance. (C) Plotting the absolute impedance as a function of cell conductance reveals curves that are almost linear at low values of conductances. These curves, however, then peak at higher conductance levels. These peaks are shifted to higher conductances when examining the curves at higher frequencies. (D) The conductance at maximal impedance (peaks at C) depends on Re and on the frequency. The red curve is the maximal possible value of the conductance at the peak impedance, regardless the value of Re.

Equation 3 shows that when G is relatively small (compared to w*C) there is a semi-linear relationship between |*Z(f)*| and G, which depends on the product of Re and G in the middle term. Indeed squaring Equation 4 and linearizing it with respect to G (i.e., G ≪ w*C) shows this linear relationship.

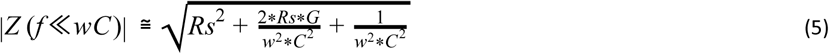

When Rs is large, linear relationship holds for both | Z(f)|^2^ or |Z(f)|. Hence, the series resistor ‘amplifies’ the cell conductance, allowing to estimate it from the electrode-cell impedance. However, the range of linearity is constrained by the cell conductance. When the cell conductance increases, the curves start to saturate until they reach a maximal impedance value (*G(Z*_*max*_)) as shown in Figure 2C. These peak values depend on the frequency, capacitance and Rs, as shown in Equation 6 (Fig. 2D).

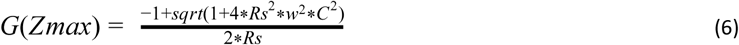

The higher is the series resistance is, the higher is *G(Z*_*max*_). For typical point neurons with a total capacitance of 1.5E-10 F, recorded with a series resistance of 100 MΩ, *G(Z*_*max*_) at 300 Hz is about 250 nS (dashed line in Fig. 2D), correspoding to an input resistance of about 3.3 MΩ, much smaller than the typical input resistance of most neurons (∼20-200MΩ, blue dashed lines for an input conductance of 1/50MΩ in Figs. 2C and D). This conductance value is much larger than the typical change in conductance (10-50 nS) that is measured during sensory stimulation (Wehr and Zador, 2003; Priebe and Ferster, 2005; Heiss et al., 2008). Hence, a steady increase in total impedance with increasing conductance can be theoretically measured until this value. For such a cell, changes in cell impedance near its input resistance for >100 Hz are clearly almost linear (Fig. 2C). Higher electrode resistance (Rs) pushes *G(Z*_*max*_) towards higher values. However, *G(Z*_*max*_) rapidly saturates, so it is never higher than *W*C* (red dash line in Fig. 1D, the limit for G(Z_max_)) with increasing R in Equation 6). For Rs of about 100 MΩ, the *G(Z*_*max*_) is about 95% of *WC.*

Altogether, this analysis suggests that it is possible to measure cell’s conductance when injecting a high-frequency sinusoidal current. However, an important trade-off influences the total conductance measurement. As the frequency of the injected current increases, so does the range where conductances can effectively be measured. Yet, the sensitivity of the measurement decreases as the frequency increases (see the positive slope in Figure 2C).

### AC analysis of a simulated passive point neuron receiving variable synaptic inputs

The major challenge in the next step is to measure the total conductance (leak conductance + synaptic conductances) of a neuron during a current clamp recording in a single trial. We addressed this by simulations of a point neuron. Above we saw that the total impedance of the cell and the electrode are nearly linearly related to the total conductance of the cell itself (*g*_*l*_*+g*_*e*_*+g*_*i*_). We sought to determine if we can estimate changes in conductance of a simulated point neuron under current clamp in a single trial by injecting a high-frequency sinusoidal current. We repeated the simulation that is shown in Figure 1, but this time included a recording electrode (Rs = 66 MΩ). In addition, we applied a step current injection in the early part of the train, to find how the step current affects the measurements. In theory, such step current should not be faulted with conductance change. We also simulated a rapid change in synaptic conductances towards the end of the trial to find how fast we can track rapid changes in conductance.

In our simulations we injected a sinusoidal current (701 Hz, 400 pA amplitude) into the model cell which receives excitatory and inhibitory synaptic inputs (Figure 3A) under two conditions. In the first case, the electrode resistance was set to zero (*Rs* = 0 M*Ω*, Fig. 3B) whereas in the second case we set it to 66 MΩ (Fig. 3E). No other parameters were changed. The voltage fluctuations were considerably larger in the second case, as expected from the large drop of the voltage across Rs due to its ohmic character, which is typically subtracted in current clamp recordings under ‘bridge mode’. Note that the sinusoidal current induced only tiny fluctuations in Vm (Fig. 3B) and thus it was never too far from its natural trajectory (if no current at all would be injected). In the simulation we also injected a current pulse at the early part of the trace to investigate how such this pulse affects the conductance measurement. To analyze the impedance of the cell at the frequency of the injected current, we first band-passed the voltage at that frequency (701 ± 1 *Hz*) and used the Hilbert transform in order to extract the modulation envelope of the band-passed voltage (using ‘envelope.m’ in Matlab). In the first case (Rs = 0 MΩ) the bandpassed signal was small and exhibited no clear modulation (Fig. 3D). Inspection of the bandpassed voltage in the second case (Rs = 66 MΩ) shows a small modulation (Fig. 3G), after dividing it by the current amplitude to obtain MΩ units. The shape of the envelope is remarkably similar to the total conductance of the cell shown in Figure 1A. This modulation is predicted from the theoretical analysis that we presented above (Fig. 2B,C). Hence, our analysis shows that the response of the cell to the sinusoidal current, when it is recorded with a series resistor, provides an envelope signal that tracks the total conductance of the cell, and that it can be extracted in a single trial.

**Figure 3:**
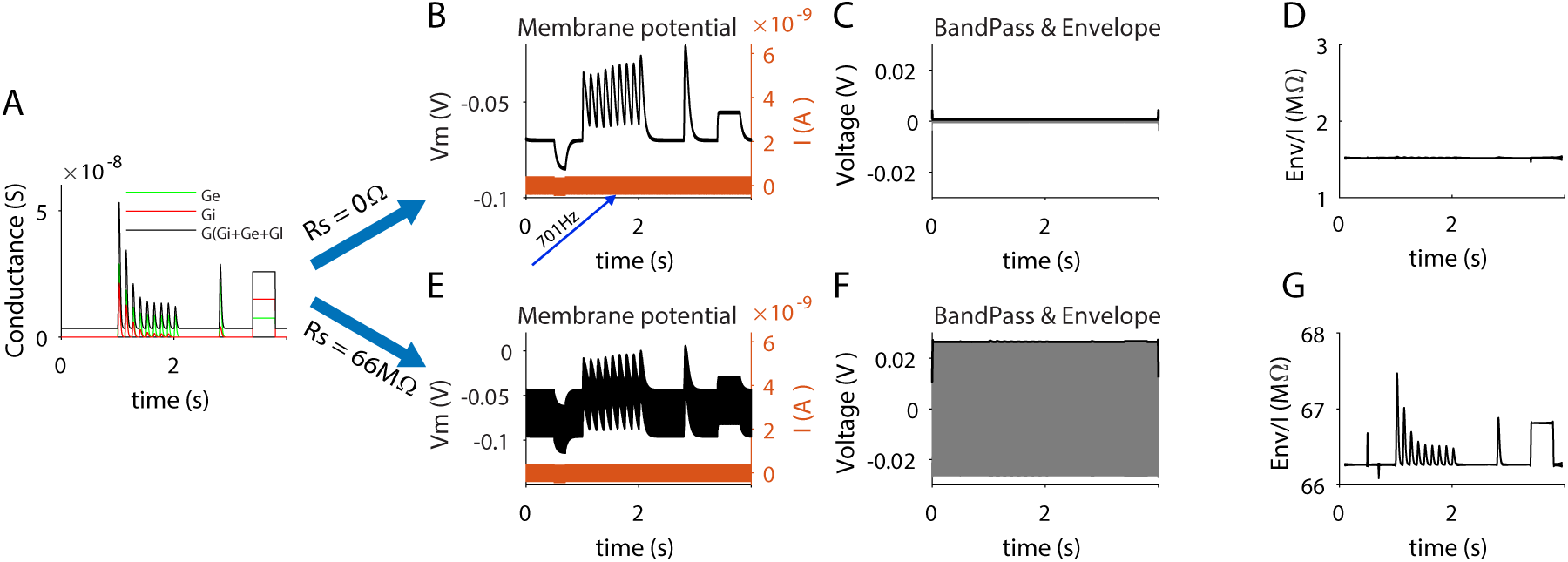
Simulation of a point neuron and recovery of the cell’s impedance at high frequency. (A) Input excitatory and inhibitory conductances of the simulated cell. (B) The membrane potential response of the cell to the synaptic inputs and injection of a sinusoidal current. Square current pulse was also injected at the early part of the trace. (C) The voltage was filtered (bandpassed at 701 ±1 *Hz*) and its Hilbert envelope is shown with the thick black line. (D) The envelope alone shows a small signal, with barely any modulation except for some small transient artifacts. (E) The same as B-D but this time when the series resistance was set to 66MΩ. In this case, the Hilbert envelope (G) is remarkably similar to the input total conductance which we set and aimed to reveal (compare the black curve in A to G).

### Extraction of excitatory and inhibitory inputs from the high-frequency voltage envelope

The analysis above shows that the impedance at the injected frequency provides a signal that is proportional to the total conductance of the cell when it is recorded in series with a resistor (i.e., the pipette resistance together with the access resistance). The next step is to convert the impedance signal to the cell’s total conductance, after subtraction of series resistance. Once this step is accomplished we can dissect the total conductance into the excitatory and inhibitory conductances, based on Equation 2 above.

Subtraction of Rs is easy if one knows exactly its value. However, in real experiments the series resistance of the electrode might change from one trial to the next due to membrane resealing and other factors. Incorrect subtraction of the series resistance using Equations 5 above and 7 below, can greatly affect the estimation of Ge and Gi. Therefore we developed an optimization process which aims at obtaining the correct electrode resistance. In this process the data is fitted with Equation 2 while minimizing two conditions which are described below. The only time-dependent input of this optimization process is the voltage from which we obtain also its impedance-envelope curve. We use three fixed constants (the reversal potential of excitation and inhibition, 0 and −0.08 V, and a measured value of total capacitance). We also describe an additional constant below. The flowchart of the process is shown in Figure 4.

**Figure 4:**
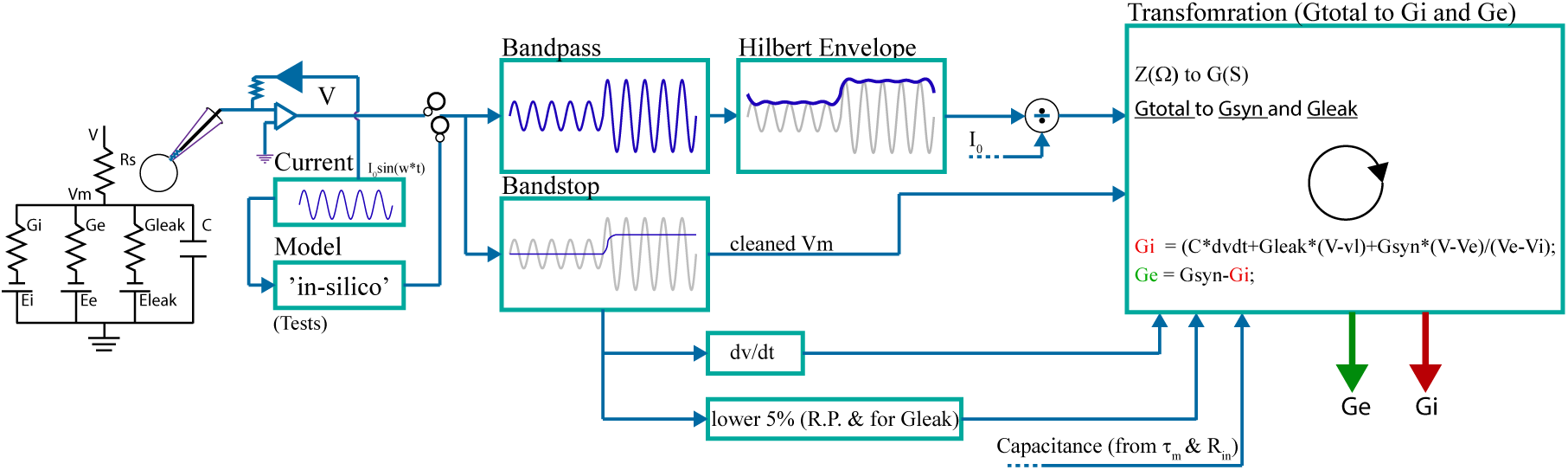
Flowchart of the method for estimating the excitatory and inhibitory synaptic conductances. Sinusoidal current is injected into a neuron. The voltage is first filtered at the frequency of the injected current and the envelope is computed before dividing it to the amplitude of the injected current to scale it to impedance units. The sinusoidal oscillations are also filtered out and the slope of the ‘cleaned voltage’ is computed. These two inputs are then fed into an optimization process which fits the two with the equation described in the right box. During the optimization process the series resistance is altered until the two conditions described in the text are met.

Equation 2 was used to calculate the excitatory and inhibitory conductances. We therefore filtered out from the membrane potential any component at the frequency of the injected sinusoidal current (‘cleaned voltage’). Before using Equation 2 we first need to transform the impedance-envelope curve to the cell’s total conductance. Although in the simulations as well as in real experiments Re can be measured, we decided not to measure it but rather evaluate it automatically using the optimization process as its value can change across trials. A proper subtraction of the electrode resistance from the total impedance is crucial as we discuss below.

Although the envelope has a very similar shape to that of the conductance curve (Fig. 3A and G), we note that the relative depth modulation of the two curves is different. The depth of the impedance-envelope relative to the electrode resistance (Rs = 66 MΩ) is smaller, i.e., its baseline is higher. Hence, subtraction of the electrode resistance will not provide a curve that has the same baseline and depth as that of the real total conductance of the cell. This is expected from Equation 4, where the second and third terms contribute additional DC level to the envelope, beyond the electrode resistance. The subtraction of the electrode resistance and thus a correct evaluation of the leak conductance is done under the optimization process as we describe below. In this process the only variable that will be altered is the series resistance (Re).

First we transform the impedance-envelope curve to a conductance curve. This transformation is part of the optimization process. The relationships between |Z(f,t)| and G(t) (from equation 4 above, after neglecting the last term when 2*pi*f*c ≫; g) can be rewritten as follows:

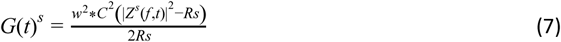

Using this equation, we transform the impedance to conductance. We marked G by uppercase s to note that this vector will be scaled during the optimization process when changing Rs. We found empirically, however, that the impedance (Z^s^ (f,t)) used here needs to be transformed from the measured impedance by the following operations (Eq. 8 is calculated before 7 during the optimization process):

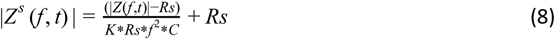

The |Z(f)| is simply obtained from the impedance-envelope curve at the frequency of the injected sine wave current (Fig. 3G) and the constant K was measured to be ∼1/1400, but its exact value is not crucial for finding E and I in an accurate manner. In equations 7 and 8 above, Rs will be changed during the optimization process as described below in order to find the exact Rs value that fulfills the optimization rules below.

After transforming the impedance envelope to the initially-guessed conductance, we sought to find the exact value of Rs that provides the best estimation of Gl, Ge(t) and Gi(t). When fitting the data with Equation 2 it is important to find the resting potential and leak conductance. This was done by finding the mean of the lower 5^th^ percentile of the total conductance (where no stimulation was given). The corresponding membrane potential values for this 5th percentile conductance were used to calculate the mean resting potential. If the electrode resistance is not well-adjusted, the total conductance curves for Ge and Gi might be different than the actual ones. Such deviations will lead to off-scaled Gleak and excitatory and inhibitory conductances, leading for example to either estimated negative excitatory or negative inhibitory conductances when both are in fact positive. Although the sum of these three conductances is constant at a particular time point (to match the total conductance curve that was transformed from the envelope-impedance curve), they can have different signs and therefore their absolute values can be much larger than the actual conductances. Our approach was to run an optimization loop when fitting the voltage (using Equations 8, 7 and 2 in this order) such that the estimated excitatory and inhibitory conductances obey together 4 optimization rules while changing Rs:

1. For the entire trial we minimized the *absolute* sum of *negative* excitatory and inhibitory conductances during times in which a positive change in envelope - transformed synaptic conductance (Ge(t)+Gi(t)) was found (minimizing abs(Ge<0) + abs(Gi<0) when Gtotal>0).
2. For the entire trial we minimized the sum of *positive* excitatory and inhibitory conductances during times in which a negative change in envelope - transformed synaptic conductance (Ge(t)+Gi(t)) was found (minimizing Ge>0 and Gi>0 when Gtotal<0).
3. For the entire trial we minimized mean(Ge(t)^2^+Gi(t)^2^)
4. As an output we used: value(1)+value(2)+value(3).

The logic behind the first rule is as follows: our assumption when measuring the leak conductance is that it is the minimal conductance during a trial, occurring when the membrane potential is at its resting potential. Therefor negative Ge or Gi are not possible in this case. However, if our estimation of the leak conductance is wrong, due to wrong subtraction of the electrode resistance, negative Ge or Gi can be measured. Yet, we added the second term since it is possible that in other cases tonic synaptic activity exists and therefore the leak conductance is ‘contaminated’. In such a case the change in total conductance can be negative and therefor we also want to minimize for these times positive change in conductance. The third rule helps also as it aimed in finding the solution with minimal conductance change.

For a given single trial voltage trace, the iterative optimization function (Matlab function ‘fminsearch.m’ or its modified version ‘fminsearchbnd.m’) was fed with the outcome of a function that calculates the excitatory and inhibitory conductances (based on Equations 8, 7 and 2 above). The output of the called function of ‘fminsearchbnd.m’ is the summed values of the three above rules as shown in (4) above, calculated as a single number for the entire trial. Again, the only variable that was changed during this optimization process was the electrode resistance (Rs). We limited the range of the search to about ±20MΩ of the initial guessed value of Rs, so as long as the true value of Rs is within this range the process converges to almost the exact value of Rs (see also below).

The results of this measurement for a simulated point neuron are shown in Figure 5. Similar to the previous figures above, we simulated the response of the cell to adapting excitatory and inhibitory inputs, followed with a step change in these conductances (Fig. 5A), while injecting sinusoidal current at 701 Hz (Fig. 5B). Two vectors were extracted from the membrane potential of the simulated cell: 1) The ‘cleaned Vm’ following filtering out any 701 ±0.1 Hz components (Fig. 5C and 2) The Hilbert envelope of the voltage after dividing it by the amplitude of the injected sinusoidal current (Fig. 5D). Excitatory and inhibitory conductances were extracted using the optimization algorithm mentioned above, assuming that the reversal potential of excitation and inhibition are known (−0.08 and 0 V) and that the total capacitance is also known (Fig. 5E). Re, measured by the optimization process was nearly the same as was set in the simulation (66.27MΩ compared to 66MΩ). Our computations revealed very similar excitatory and inhibitory conductances to those used to simulate the response of the cell (compare Fig 5E to A). Note that our algorithm also revealed the precise timing between the inputs – in our simulation we imposed a delay of 3 ms between excitation and inhibition (inset in Fig. 3A, depicting the response to the first stimulus in the train). This delay was also revealed in the extracted conductances (inset in Fig. 5E). Hence, the entire approach described above allows measuring excitatory and inhibitory conductances in a single trial from a single neuron with high temporal precision.

**Figure 5:**
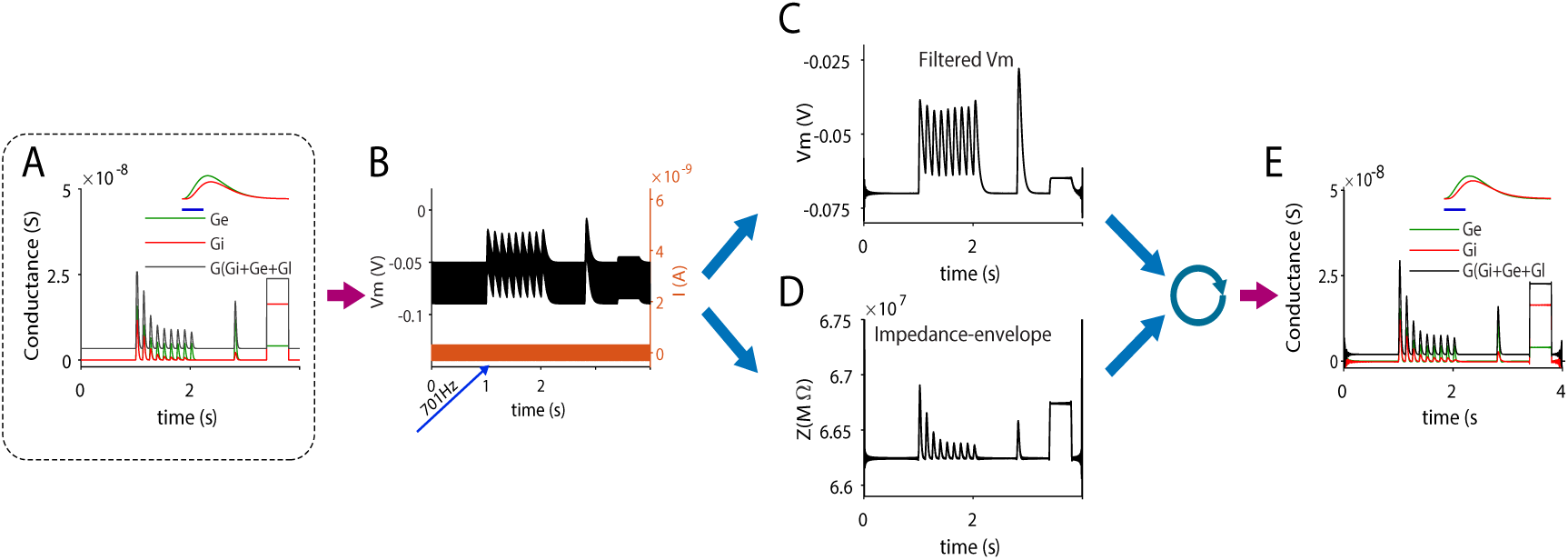
Simulation of the method for the estimation of excitatory and inhibitory conductances in a single trial during current clamp recording of a point neuron. (A) The synaptic inputs of a simulated point neuron. (B) The membrane potential of the cell due to the synaptic inputs in A as well as the 701 Hz sinusoidal current injection. (C) Sinusoidal component in B is filtered out to obtain the cleaned voltage. (D) The Hilbert envelope of the voltage in B. (E) Estimated excitatory and inhibitory conductances as obtained following the optimization process (see text for more details). Note the similarity of these estimated conductances to those set as inputs in the simulation (Compare A to E). Insets in A and E show that the temporal relationships between excitation and inhibition were also revealed.

Finally, we asked if our approach can be used to reveal the underlying excitatory and inhibitory conductances of a cortical neuron embedded in an active network where it receives excitatory and inhibitory inputs. Therefore we used a simulation of a cortical network at a balanced asynchronous state (Renart et al., 2010) to obtain the excitatory and inhibitory synaptic inputs of a single cell (kindly provided by Dr. Michael Okun, University of Leicester). We used these conductances in an additional simulation of a single cell, in which we injected a sinusoidal current (750 Hz) via a 100 MΩ electrode (Fig. 6). The voltage response of the electrode + cell exhibits strong sinusoidal modulation at the current frequency (Fig. 6A), which was mostly due to the drop of voltage across the recording electrode (Rs). We filtered out the 750 Hz component of the total voltage (Fig. 6B, black trace), allowing us to observe the fluctuations of the cell membrane potential due to the synaptic inputs, as if no current was injected. Indeed, this trace is very similar to the one obtained without current injection and when setting the electrode resistance to zero (Fig 6B, blue trace). Next, we used the optimization process described above to reveal the excitatory and inhibitory inputs of the simulated cell. The electrode resistance was estimated following this process to be 102.27 MΩ, slightly higher than the 100 MΩ which was used in the simulation. The extracted excitatory and inhibitory synaptic inputs were very similar to those used as inputs (Fig. 6C-F). Note, however, that for both inputs the extracted conductances are more negative than expected. This is simply because the leak conductance was estimated from the 5^th^ percentile of the total conductance of the cell, but since synaptic activity persisted throughout the trace, the leak conductance reflects a mixture of the true leak conductance and some baseline synaptic activity. In summary, this simulation shows that the membrane potential dynamics of a neuron can be obtained in a single trial while estimating its underlying excitatory and inhibitory synaptic conductances.

**Figure 6:**
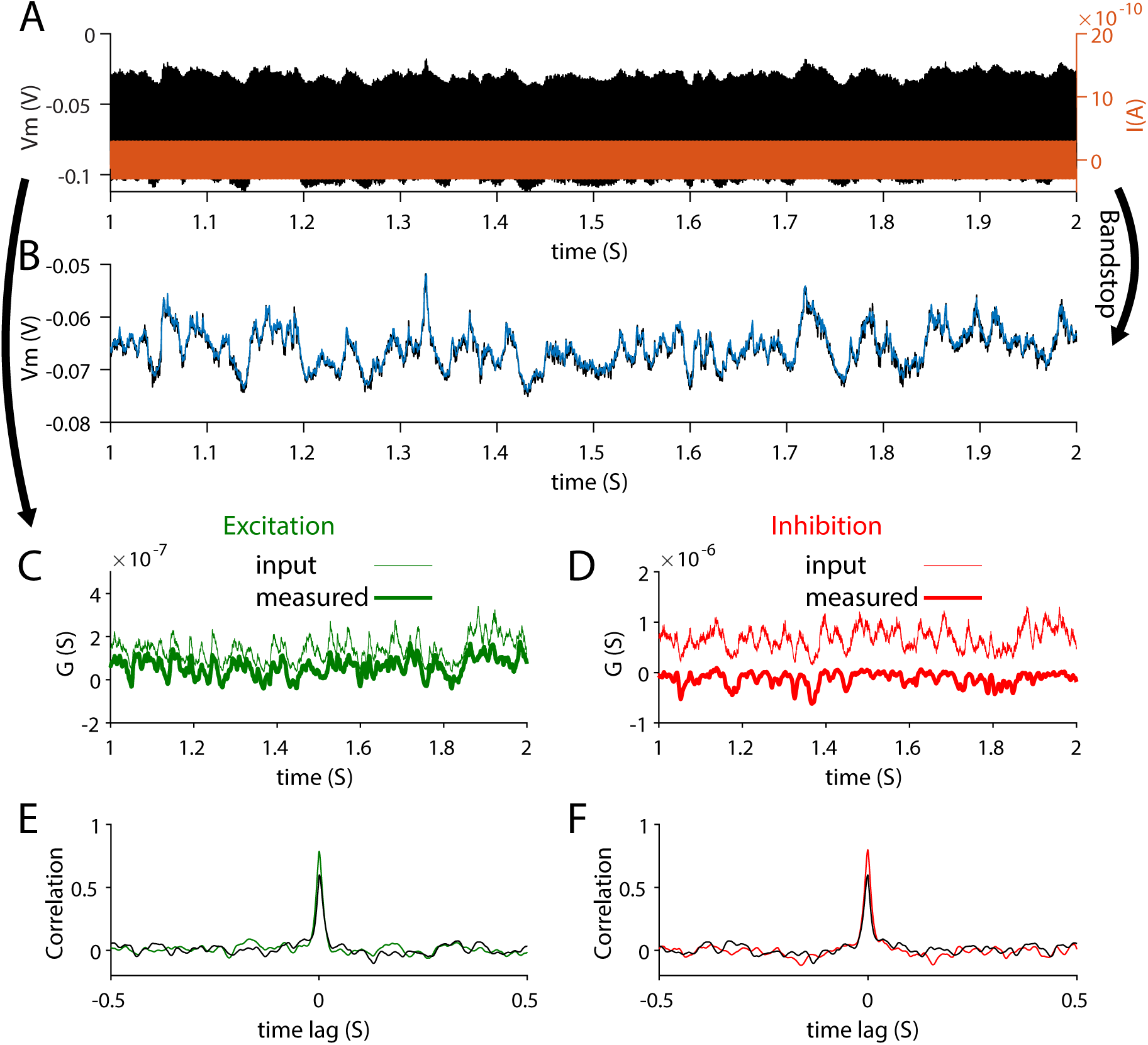
Estimation of excitatory and inhibitory conductances of a neuron embedded in a network that exhibit balanced asynchronous state. (A) Excitatory and inhibitory conductances were used to simulate the response of a point neuron recorded with a 100MΩ pipette while injected with 750 Hz sinosiodal current. (B) The voltage in A was ‘cleaned’ from the sinusoidal component (black trace) and it is displayed with the voltage response of the cell when no current was injected and in absent of series resistance of an electrode. (C-D) Estimated excitatory and inhibitory conductances (thick traces) superimposed with the imposed conductances of the simulated cell (thin traces). (E-F) Color lines describe the cross-correlations between the imposed and estimated conductances and the black lines are the correlations between imposed (real) Ge and Gi, which are lower than the real-measured correlations for each input.

## Discussion

We describe a novel method that in principle enables estimating the excitatory and inhibitory synaptic conductances of a neuron in a single trial with high temporal resolution while tracking the trajectory of the membrane potential when recorded under current clamp mode. The work described above is theoretical and lays the foundations for future experimental work.

Our method is based on AC analysis of RC circuits. We show that theoretically, the response of the voltage to sinusoidal current input across the Re-RC circuit is amplified by the conductance of the cell, enabling us to reveal changes in cell’s conductance. This amplification is revealed when computing the Hilbert envelope of the voltage. Changes in cell conductance were then fed into an optimization process that revealed the excitatory and inhibitory conductances at each time point in a single trial.

We demonstrated the method in two cases. In one we showed that we can reveal the timing and magnitude of discrete adapting excitatory and inhibitory synaptic inputs. In another example, we used our method to reveal these inputs during an asynchronous balanced cortical state. Importantly, these estimations were obtained at a single trial and allow obtaining the natural dynamics of the membrane potential by filtering out the sinusoidal response to the injected current.

### Comparisons with other methods

#### Measurement of average excitatory and inhibitory conductances of single cells

Excitatory and inhibitory synaptic conductances of a single cell were measured both under voltage clamp or current clamp recordings, focusing *in-vivo* on the underlying mechanisms of feature selectivity in sensory response of cortical cells and in particular on the role of inhibition in shaping the tuned sensory response of mammalian cortical neurons (Borg-Graham et al., 1998; Wehr and Zador, 2003, 2005); (Anderson et al., 2000; Tan et al., 2004; Priebe and Ferster, 2005; Wilent and Contreras, 2005; Heiss et al., 2008). Conductance measurement methods were also used to reveal the underlying excitatory and inhibitory conductances during ongoing Up and Down membrane potential fluctuations, which characterize slow wave sleep activity (Shu et al., 2003; Haider et al., 2006). The advantages and caveats of these methods were reviewed in (Monier et al., 2008). Common to these conductance measurement methods is the requirement to average the data over multiple repeats, triggered on a stereotypical event (such as the time of sensory stimulation or the rising phase of an Up state) and the need to record the activity at different holding potentials. Data obtained under different holding potentials allowed to shift the membrane potential away from the reversal potential of a given conductance and thus to increase the driving force for particular ions that flow via specific channels. In these studies, similar to our new method, the reversal potential of excitation is assumed to be around 0 mV and that of inhibition at about −80 mV. Likewise the averaged data is then fitted with the membrane potential equation to reveal the conductance of excitation and inhibition at each time point. However, these methods cannot reveal inhibition and excitation simultaneously and only estimate averaged relationships. Our proposed AC measurement method, on the other hand, allows for simultaneous measurements at a single trial.

An alternative approach for estimating the excitatory and inhibitory conductances of a single cell was demonstrated for retinal ganglion cells (Cafaro and Rieke, 2013). In this study the clamped voltage was alternated between the reversal potential of excitation and inhibition at a rate of 50 Hz and the current was measured at the end of each step. This study revealed strong correlated noise in the strength of both types of synaptic inputs. However, unlike the method proposed here, the underlying conductances are not revealed simultaneously and due to the clamping, the natural dynamics of the membrane potential is completely unavailable, preventing examining the role of intrinsic dynamics in the generation of neuronal subthreshold activity.

Theoretical approaches based on the statistical estimation of excitatory and inhibitory conductances in a single trial were proposed before. Accordingly, excitation and inhibition are revealed from current clamp recordings in which no current is injected. These approaches are based on Baysian methods which exploit multiple recorded trials (Lankarany et al., 2016). Single trial estimation of these inputs in a single trial were also proposed but lack the ability to track fast changes in these conductances (Paninski et al., 2012).

#### Paired intracellular recordings

The substantial synchrony of the synaptic inputs among nearby cortical cells (Lampl et al., 1999; Hasenstaub et al., 2005; Okun and Lampl, 2008; Malina et al., 2016) allows continuously monitoring both the excitatory and inhibitory activities in the local network. Toward this end, simultaneous recordings from nearby pairs of cortical neurons in anesthetized rats were taken, where one cell was hyperpolarized close to the reversal potential of inhibition to reveal excitatory inputs and the other cell was depolarized close to the reversal potential of excitation to reveal inhibitory inputs (Okun and Lampl, 2008). A similar approach was recently used to study the relationships between these inputs in the visual cortex of awake mice (Arroyo et al., 2018) as well as gamma activity in slices (Atallah and Scanziani, 2009). While paired recordings are powerful when examining the relationships between these inputs in the local network, such recordings do not provide definitive information about the inputs of a single cell. Moreover, interpreting changes in the correlations between these inputs across brain-states is not straightforward. For example, a reduction in the correlation between excitation, as measured in one cell, and inhibition measured in the other cell can truly suggest smaller correlation between these inputs for each cell, but it can also result from a reduction of synchrony between cells, without any change in the degree of correlation between excitation and inhibition of each cell. This caveat of paired recordings prevents us from finding, for example, if cortical activity shifts between balanced and unbalanced states (Tsodyks and Sejnowski, 1995; van Vreeswijk and Sompolinsky, 1998). Simultaneous measurement of excitatory and inhibitory conductances of a single cell across states will allow these and other questions to be addressed.

### Key principles of the new method

The starting point of this work is demonstrated in Figure 1, and it consists of the fact that excitatory and inhibitory conductances can be revealed if we can record the membrane potential and at the same time measure the total conductance of the cell driven by synaptic inputs. Hence, the challenge is to reveal the conductance of the cell with high temporal resolution. We found a method to estimate rapid changes in the cell’s total conductance. The method is based on AC analysis of RC circuit, connected in series to another resistor which represents the ohmic impedance of the recording pipette. Surprisingly, at high frequencies, the impedance of this circuit increases when the cell’s conductance gets higher. For sufficiently high frequencies, changes in the absolute impedance of this circuit are almost linearly correlated with a change in the cell’s total conductance, allowing us to calculate the total conductance of the cell over time. The final step in our method involves an optimization process in which we extract the excitatory and inhibitory conductances at each time point using the membrane potential equation. During this process we alter only one variable, the electrode resistance, minimizing the absolute sum of negative estimated excitatory and inhibitory conductances across the entire trace when the change in total conductance is positive (and vice versa when the change in total conductance is negative). This method allows estimating the excitatory and inhibitory conductances during a single trial with high temporal resolution, limited by the frequency of the injected current.

### Limitations

We show that, theoretically, increasing the frequency of the current improves the temporal precision when measuring synaptic conductances. A higher frequency also increases the dynamic range of the measured conductance. However, this comes at the expense of sensitivity, which reduces as frequency increases (i.e., the change in impedance due to conductance changes becomes smaller, see Fig. 2C). In our simulations we limited the frequency of the injected current to about 700-750 Hz. Experimentally, we calculated and tested the time constant of typical patch pipettes (due to its resistance and stray capacitance) and found that they start to attenuate the voltage above this frequency range (not shown), suggesting that it may be possible to measure conductance changes at this temporal resolution.

In whole cell patch recordings the total series resistance is composed of the pipette resistance together with the access resistance, which is usually higher than the former due to incompletely ruptured membrane. A critical experimental requirement for our method to work is the need for a stable series resistance. There are several consequences of this. First, since instability in access resistance is more common in *in vivo* recordings compared to brain slices or cultured cells, our method is likely to be more difficult to apply *in-vivo*. Second, it is possible, theoretically, to increase the dynamic range of the measured conductance by increasing the total series resistance (Fig. 2D). Hence, adding a passive resistor between the headstage and the pipette should improve the sensitivity of our method. Note that above some value which depends on cell properties and frequency, a higher electrode resistance will not improve the range of detected conductance and the sensitivity (Fig. 2D).

Other aspects that might reduce the sensitivity of our method is the presence of stray capacitances additional to that of the pipette. Biological factors that may affect the usability of our method experimentally include current escape via dendrites and the location of synaptic inputs, considering that many inputs are located at the distal parts of the dendrites. These issues, which are well-known problems also for the standard approaches, may reduce the ability to detect conductance changes and should be tested in future theoretical and experimental studies.

### Application for measurement of intrinsic conductances

Unlike other methods for conductance estimation, we can use our method when the cell is recorded at its resting potential, without any DC current injection, allowing the voltage to evolve naturally due to synaptic and intrinsic conductances. When high-frequency current is injected into the cell via the high-resistance electrode, the drop in voltage is mostly across the electrode, implying that the actual sinoisodal fluctuations of voltage across the membrane are small (Fig. 3). Hence, voltage-dependent conductances will evolve naturally, without being greatly affected by this sinusoidal current, allowing measuring changes in conductance due to intrinsic mechanisms, including voltage-dependent conductances. Such an approach can be used *in vitro* to test agonists and antagonists for various ion channels as well as to examine the effect of neuromodulators in a single trial. With some assumptions on the reversal potential of the studied channels, our approach can be used for studying more than one type of ion channel at the same time.

In summary, our theoretical study shows that synaptic and other conductances can be measured at high temporal resolution in a single trial when cells are recorded at their resting potential. More research is needed to find if this approach can be used successfully during physiological recordings from real neurons.

## Acknowledgements

I would like to thank Dr. Ana Parabucki, Hagay Famini and Daniel Müller for checking and testing some computational aspects that were used in this study as well as for their important comments on the manuscript. I thank Dr. Yonatan Katz and Michael Sokoletsly for outstanding comments on the manuscript. I thank Dr. Michael Okun for his comments on the manuscript and for providing the simulated data used in Figure 6. This work was supported by grants from the DFG-SFB 1089, 01EW1606 - DeCipher EraNet Neuron, HFSP, Israel Science Foundation (ISF 1539/17) and Minerva. I.L. is the incumbent of the Norman and Helen Asher Professorial Chair.

## References

Adesnik H (2017) Synaptic Mechanisms of Feature Coding in the Visual Cortex of Awake Mice. Neuron 95:1147–1159.e4.

Adesnik H (2018) Layer-specific excitation/inhibition balances during neuronal synchronization in the visual cortex. J Physiol 596:1639–1657.

Anderson JS, Carandini M, Ferster D (2000) Orientation tuning of input conductance, excitation, and inhibition in cat primary visual cortex. J Neurophysiol 84:909–926.

Arroyo S, Bennett C, Hestrin S (2018) Correlation of Synaptic Inputs in the Visual Cortex of Awake, Behaving Mice. Neuron 99:1289–1301.e2.

Atallah BV, Bruns W, Carandini M, Scanziani M (2012) Parvalbumin-expressing interneurons linearly transform cortical responses to visual stimuli. Neuron 73:159–170.

Atallah BV, Scanziani M (2009) Instantaneous modulation of gamma oscillation frequency by balancing excitation with inhibition. Neuron 62:566–577.

Borg-Graham LJ, Monier C, Frégnac Y (1998) Visual input evokes transient and strong shunting inhibition in visual cortical neurons. Nature 393:369–373.

Cafaro J, Rieke F (2013) Regulation of spatial selectivity by crossover inhibition. J Neurosci 33:6310–6320.

Haider B, Duque A, Hasenstaub AR, McCormick DA (2006) Neocortical network activity in vivo is generated through a dynamic balance of excitation and inhibition. J Neurosci 26:4535–4545.

Haider B, Krause MR, Duque A, Yu Y, Touryan J, Mazer JA, McCormick DA (2010) Synaptic and network mechanisms of sparse and reliable visual cortical activity during nonclassical receptive field stimulation. Neuron 65:107–121.

Hasenstaub A, Shu Y, Haider B, Kraushaar U, Duque A, McCormick DA (2005) Inhibitory postsynaptic potentials carry synchronized frequency information in active cortical networks. Neuron 47:423–435.

Heiss JE, Katz Y, Ganmor E, Lampl I (2008) Shift in the balance between excitation and inhibition during sensory adaptation of S1 neurons. J Neurosci 28:13320–13330.

Lampl I, Reichova I, Ferster D (1999) Synchronous membrane potential fluctuations in neurons of the cat visual cortex. Neuron 22:361–374.

Lankarany M, Heiss JE, Lampl I, Toyoizumi T (2016) Simultaneous Bayesian Estimation of Excitatory and Inhibitory Synaptic Conductances by Exploiting Multiple Recorded Trials. Front Comput Neurosci 10:110.

Li Y-T, Ma W-P, Li L-Y, Ibrahim LA, Wang S-Z, Tao HW (2012) Broadening of inhibitory tuning underlies contrast-dependent sharpening of orientation selectivity in mouse visual cortex. J Neurosci 32:16466–16477.

Malina KC-K, Mohar B, Rappaport AN, Lampl I (2016) Local and thalamic origins of correlated ongoing and sensory-evoked cortical activities. Nature Communications 7 Available at: http://dx.doi.org/10.1038/ncomms12740.

McCormick DA, Contreras D (2001) On the cellular and network bases of epileptic seizures. Annu Rev Physiol 63:815–846.

Monier C, Fournier J, Frégnac Y (2008) In vitro and in vivo measures of evoked excitatory and inhibitory conductance dynamics in sensory cortices. J Neurosci Methods 169:323–365.

Moore AK, Weible AP, Balmer TS, Trussell LO, Wehr M (2018) Rapid Rebalancing of Excitation and Inhibition by Cortical Circuitry. Neuron 97:1341–1355.e6.

Okun M, Lampl I (2008) Instantaneous correlation of excitation and inhibition during ongoing and sensory-evoked activities. Nat Neurosci 11:535–537.

Paninski L, Vidne M, DePasquale B, Ferreira DG (2012) Inferring synaptic inputs given a noisy voltage trace via sequential Monte Carlo methods. J Comput Neurosci 33:1–19.

Poulet JFA, Petersen CCH (2008) Internal brain state regulates membrane potential synchrony in barrel cortex of behaving mice. Nature 454:881–885.

Priebe NJ, Ferster D (2005) Direction selectivity of excitation and inhibition in simple cells of the cat primary visual cortex. Neuron 45:133–145.

Reimer J, Froudarakis E, Cadwell CR, Yatsenko D, Denfield GH, Tolias AS (2014) Pupil fluctuations track fast switching of cortical states during quiet wakefulness. Neuron 84:355–362.

Renart A, de la Rocha J, Bartho P, Hollender L, Parga N, Reyes A, Harris KD (2010) The asynchronous state in cortical circuits. Science 327:587–590.

Schevon CA, Weiss SA, McKhann G Jr, Goodman RR, Yuste R, Emerson RG, Trevelyan AJ (2012) Evidence of an inhibitory restraint of seizure activity in humans. Nat Commun 3:1060.

Scholl B, Tan AYY, Priebe NJ (2013) Strabismus disrupts binocular synaptic integration in primary visual cortex. J Neurosci 33:17108–17122.

Shu Y, Hasenstaub A, Badoual M, Bal T, McCormick DA (2003) Barrages of synaptic activity control the gain and sensitivity of cortical neurons. J Neurosci 23:10388–10401.

Stringer C, Pachitariu M, Steinmetz N, Reddy CB, Carandini M, Harris KD (2019) Spontaneous behaviors drive multidimensional, brainwide activity. Science 364:255.

Tan AYY, Zhang LI, Merzenich MM, Schreiner CE (2004) Tone-Evoked Excitatory and Inhibitory Synaptic Conductances of Primary Auditory Cortex Neurons. Journal of Neurophysiology 92:630–643 Available at: http://dx.doi.org/10.1152/jn.01020.2003.

Tsodyks MV, Markram H (1997) The neural code between neocortical pyramidal neurons depends on neurotransmitter release probability. Proc Natl Acad Sci U S A 94:719–723.

Tsodyks MV, Sejnowski T (1995) Rapid state switching in balanced cortical network models. Network: Computation in Neural Systems 6:111–124 Available at: http://dx.doi.org/10.1088/0954-898x_6_2_001.

van Vreeswijk C, Sompolinsky H (1998) Chaotic Balanced State in a Model of Cortical Circuits. Neural Computation 10:1321–1371 Available at: http://dx.doi.org/10.1162/089976698300017214.

Vinck M, Batista-Brito R, Knoblich U, Cardin JA (2015) Arousal and locomotion make distinct contributions to cortical activity patterns and visual encoding. Neuron 86:740–754.

Wehr M, Zador AM (2003) Balanced inhibition underlies tuning and sharpens spike timing in auditory cortex. Nature 426:442–446.

Wehr M, Zador AM (2005) Synaptic mechanisms of forward suppression in rat auditory cortex. Neuron 47:437–445.

Wilent WB, Contreras D (2005) Dynamics of excitation and inhibition underlying stimulus selectivity in rat somatosensory cortex. Nat Neurosci 8:1364–1370.

Wu GK, Arbuckle R, Liu B-H, Tao HW, Zhang LI (2008) Lateral sharpening of cortical frequency tuning by approximately balanced inhibition. Neuron 58:132–143.

Zhou M, Liang F, Xiong XR, Li L, Li H, Xiao Z, Tao HW, Zhang LI (2014) Scaling down of balanced excitation and inhibition by active behavioral states in auditory cortex. Nat Neurosci 17:841–850.

